# Vascular endothelial growth factor receptors 1 and 3 mediate placental trophoblast leptin production in preeclampsia, inducing vascular dysfunction

**DOI:** 10.1101/2025.05.27.656489

**Authors:** Mona Elgazzaz, Safia Ogbi, Desmond Moronge, Elisabeth Mellott, Gibson Cooper, Kristin Backer, Joanna Hitchings, Luis Valesquez Zarate, Sravankumar Kavuri, Tae Jin Lee, Daria Ilatovskaya, Padmashree C. Woodham, James Maher, Brian H. Annex, Jessica L Faulkner

## Abstract

Heightened soluble FMS-like tyrosine kinase-1 (sFlt-1) levels is a hallmark of preeclampsia patients and induces a state of angiogenic imbalance by sequestering free vascular endothelial growth factor (VEGF) and placental growth factor (PlGF). The receptors for VEGF and PlGF, membrane-bound VEGFR, are expressed in placental trophoblast cells, but their functions are largely unknown. Placenta production of leptin significantly increases in preeclampsia, and we recently showed leptin induces placental and vascular endothelial dysfunction in pregnancy. We hypothesized that there is a mechanistic link in which inappropriately high sFlt-1 in preeclampsia leads to an increase in trophoblast leptin production. We treated human placental explants and trophoblast cells with sFlt-1 and show an increase in leptin peptide production, which is ablated by coadministration with either VEGF or placental growth factor (PLGF). We further demonstrate that VEGFR1 and 3, not R2, expressions are predominant in human trophoblasts and that reducing activation of these receptors mediates trophoblast leptin production. In pregnant mice, we show that sFlt-1 infusion induces vascular endothelial dysfunction in association with significantly elevated plasma leptin levels. In pregnant sFlt-1-infused mice treatment with leptin receptor antagonist significantly ablated vascular endothelial dysfunction. Collectively, these data indicate that angiogenic imbalance in preeclampsia impacts placental trophoblast endocrine function by suppressing VEGFR1 and 3 activation, resulting in leptin overproduction. Furthermore, sFlt-1 induces vascular endothelial dysfunction in mice dependent on leptin receptor activation.

## Introduction

Preeclampsia is a leading cause of maternal mortality causing more than 70,000 maternal and half a million fetal deaths each year worldwide.^1–4^ Preeclampsia is an increasingly prevalent hypertensive disorder of pregnancy presenting with hypertension in mid-late gestation and associated endothelial dysfunction.^5^ Soluble FMS-like tyrosine kinase 1 (sFlt-1), is produced from the placenta trophoblasts at elevated levels in preeclampsia and is strongly associated with adverse outcomes.^6–9^ sFlt-1 is a soluble form of vascular growth factor receptor 1 (VEGFR1) which lacks the transmembrane domain and thus is unable to activate cellular signaling, but retains the ability to sequester VEGFR1 ligands including VEGF and placental growth factor (PlGF). ^10,11^ In this capacity sFlt-1 downregulates VEGFR activity and is a major component of the anti-angiogenic milieu that presents in preeclampsia patients. The implications of high sFlt-1 on vascular endothelial cells, centrally mediated through suppression of VEGFR2 activity, are well-characterized in preeclampsia.^12^ However, despite that VEGFRs are prominently expressed on trophoblast cells, the function of these receptors and the potential detrimental effect of sFlt-1 on trophoblast VEGFR signaling is largely unknown.

Similar to sFlt-1, numerous clinical studies show a strong association of elevated plasma leptin levels in preeclampsia patients, independent of body mass index.^13–24^ In nonpregnant individuals adipose tissue is the major source of leptin production and its levels are heavily governed by adipose tissue mass. However, in pregnancy, placental trophoblasts produce high levels of leptin, and several studies indicate that preeclampsia pathologically increases trophoblast leptin production.^25^ Work by our group and others shows that infusion of leptin in pregnant rodents induces key features of preeclampsia including endothelial dysfunction, hypertension and fatal growth restriction.^26–28^ However, placental sFlt-1 levels do not increase in response to leptin infusion, despite the development of other key characteristics of preeclampsia.^26^ We hypothesized that sFlt-1 mediates increases in placental trophoblast leptin production in preeclampsia placenta, leading to adverse vascular outcomes.

## Methods

### Human Placental Explants

All studies involving human tissues were approved by the Augusta University Institutional Review Board. Placental tissue was collected within 6 hours of delivery from patients delivering at Wellstar Medical College of Georgia Health Hospital in Augusta, GA. Healthy patients were singleton pregnancies delivered full term (>37 weeks) without diagnosis of preeclampsia, gestational diabetes, chronic hypertension of pregnancy, multiple births, fetal growth restriction or other notable diagnoses. Preeclampsia was diagnosed by American College of Obstetrics and Gynecology criteria and singleton pregnancies were collected. Placental explants were harvested from the maternal side proximal to the umbilical cord insertion site and rinsed with (PBS+1%EDTA+20%Pen/Strep).^29^ Explants were cultured in a 6-well plate on Matrigel© in F-12K+1%Penn/Strep+10%BSA and treated with either sFlt-1 (1ng/mL, 72 h; recombinant human VEGFR1/Flt-1 Fc chimera protein #321-FL-050/CF, R&D systems, USA), VEGF (10 ng/mL, 24 h, recombinant human VEGF165 protein #293-VE, R&D systems, USA), PlGF (5 ng/mL, 24 h, recombinant human PlGF protein #264-PGB/CF, R&D systems, USA), a combination, or a saline vehicle for 72 hours at 37°C in 95% O_2_/5% CO_2_. Following incubation, media were collected for analysis.

### Cell Culture of human trophoblast cells BeWo

For cell culture experiments, Human BeWo trophoblasts (ATCC©, USA) at 0.5-2×10^5^ cells/well (70% confluence) in OptiMEM were serum-starved (1%) 24h prior to experiment. Cells were then incubated with sFlt-1, VEGF, PlGF, or combination in the same concentrations as placental explants for 72 hours. Cells and media were harvested for analysis.

### Quantitative PCR

Tissues and cells were homogenized using TRIzol and RNA was isolated with RNA mini kit (#12183018A, Invitrogen, USA). cDNA was then obtained using reverse transcription (ThermoFisher, USA) followed by qRT-PCR with Sybr Green (Applied Biosystems, USA). Primer sequence used are in Table S1. Ct values were normalized to 18s expression (ΔCt) followed by normalization to control groups (ΔΔCt) and calculation of relative gene expression (2-ΔΔCt) as published previously^26^ and in accordance with Applied Biosystems guidelines for expression analysis (Applied Biosystems User Bulletin, no. 2, 1997).

### ELISA

PLGF, VEGF and Leptin were measured in culture media according to the manufacturer’s protocols (PLGF: #DPG00, R&D Systems; VEGF: #DVR100, R&D Systems; Leptin: RAB0333-1KT, Millipore Sigma). In mice, plasma Leptin (# EZML-82K, Millipore Sigma, Danvers, MA) was quantified using enzyme-linked immunosorbent assays (ELISA) following the manufacturer’s protocols.

### Western Blot

BeWo or HUVEC cell lysates were prepared, and proteins separated by SDS-PAGE (Bio-Rad). Following separation, proteins were transferred to polyvinylidene difluoride (PVDF) membranes (Immun-Blot, Bio-Rad). Membranes were then blocked at room temperature (22 °C) in Tris-buffered saline containing 0.05% Tween-20 (TTBS) and 5% (w/v) non-fat dry milk. Blocked membranes were incubated overnight at 4 °C with primary antibodies diluted 1:1000 in blocking buffer. The following primary antibodies were used: anti-phospho-VEGFR1 (pVEGFR1; Sigma, Cat# SAB4504006), anti-VEGFR1 (Abcam, Cat# ab32152), anti-phospho-VEGFR2 (pVEGFR2; R&D Systems,Cat# AF1766), anti-VEGFR2 (Abcam, Cat# ab39638), anti-phospho-VEGFR3 (pVEGFR3; MyBioSource, Cat# MBS9612744), and anti-VEGFR3 (Invitrogen, Cat# PA5-109730), anti-phospho-ERK1/2 (pERK1/2; Cell Signaling Technology, Cat# 4370), and anti-ERK1/2 (Cell Signaling Technology, Cat# 4695). HSP90 (BD Transduction Laboratories, Cat# 610419) was used as a loading control. VEGFR antibodies were validated by negative control. Membranes were washed and incubated for 1 hour at room temperature with horseradish peroxidase (HRP)-conjugated secondary antibodies diluted 1:1000 in blocking buffer including Anti-rabbit IgG-HRP (Cell Signaling Technology, Cat# 7074) or anti-mouse IgG-HRP (Cell Signaling Technology, Cat# 7076). Protein bands were visualized using enhanced chemiluminescence (ECL) substrate (Cytiva, Cat# RPN2236) and detected using the FluorChem E imaging system (ProteinSimple). Molecular weight markers (Bio-Rad) were run in parallel to confirm the identity and size of the detected bands. Protein loading was normalized by probing for heat shock protein 90 (HSP90; BD Transduction Laboratories, Cat# 610419), used at a dilution of 1:1000. Data were quantified using Image J software.

### Immunofluorescence and Confocal Microscopy

BeWo cells were seeded on 4 well-chamber slide (Nunc™Lab-Tek™ II #154526PK, ThermoFisher, USA). After 24 h, the cells were treated with sFlt-1 (1ng/mL, 72 h) or PBS. Cells were then rinsed with 1X PBS, and fixed with 4% PFA at room temperature (RT) for 30 min. The cells were 5X washed twice with 1X PBS at RT. Further, the cells were permeabilized in buffer (0.1% Triton X-100 detergent in 1X PBS) at RT for 20 min then washed with 1X PBS at RT for 3 times. Cells were then blocked for 1 hour at room temperature (2% BSA in 1X PBS). The blocking buffer was replaced with Leptin primary antibody (1:100, # PA1-052, ThermoFisher, USA) diluted1% BSA in PBS at 4 °C, overnight. The cells were further washed 3 times with PBS at RT. The cells were treated with combined Fluorescein Phalloidin-FITC (1:250 #F432, ThermoFisher, USA) and the secondary antibody (#A-31573,ThermoFisher, USA) diluted in 1% BSA in 1X PBS for 1 hour at room temperature in the dark. Cells were then washed, and mounting media DAPI (ProLong™ Diamond Antifade Mountant with DAPI, # 359-460, ThermoFisher, USA) was added for visualization under the confocal microscope (FV3000, FluoView, Olympus, Japan). Images were taken at 40X magnification.

### In vitro VEGFR knockdown

BeWo cells were transfected with siRNA targeting VEGFR1 (siRNA ID: 144806; Catalog #AM16708; Invitrogen, Thermo Fisher Scientific), VEGFR2(siRNA ID: s535305; Catalog # 4392420; Invitrogen, Thermo Fisher Scientific) and VEGFR3 (siRNA ID: s5294; Catalog #4392420; Invitrogen, Thermo Fisher Scientific), or a non-targeting control siRNA: Invitrogen, Thermo Fisher Scientific), using Lipofectamine RNAiMAX Transfection Reagent (9 µL per well; Catalog #13778075; Invitrogen, Thermo Fisher Scientific) in serum-free Opti-MEM Reduced Serum Medium (Catalog #11058021; Invitrogen, Thermo Fisher Scientific). Transfection complexes were prepared by incubating RNAiMAX with siRNA (final concentration: 10 nM) in Opti-MEM for 5 minutes at room temperature. The complexes were then added to each well of a 6-well plate containing BeWo cells and incubated for 8 hours. After incubation, the medium was replaced with complete growth medium, and the cells were cultured for an additional 72 hours in serum free media. Gene silencing was confirmed 48 hours post-transfection by Western blot analysis. The media were then collected for leptin concentration analysis via ELISA.

### RNA sequencing of trophoblast BeWo cells

BeWo cells were treated with sFlt-1 or vehicle as above and harvested and RNA isolated as described above. Raw sequencing reads in FASTQ format were obtained from the sequencing provider (Novogene©, Sacramento, CA), where adapter trimming was performed. Transcriptome alignment was conducted using STAR (v2.7) with the GRCh38 human reference genome for both mapping and annotation. Default parameters were applied. Feature counting was performed using subread (v2.0.3) to generate gene-level count matrices based on the aligned reads. Differential expression analysis was conducted using DESeq2 in R, following standard normalization and statistical modeling procedures. Genes with a p-value < 0.05 were considered significantly differentially expressed. All computational analyses were performed in an R-based environment, and statistical significance was determined using appropriate multiple-testing corrections where applicable.

### Experimental Animals

BALB/c mice female (10-12 weeks, Jackson Lab, USA) were bred with age-matched BALB/c males to generate timed-pregnancy. Gestation day (GD)1 was determined via the detection of a vaginal plug. All protocols were approved by the Institutional Animal Care and Use Committee (IACUC Protocol # 2011-0108) of Augusta University. Mice were housed in a temperature-controlled facility (∼24 °C) under 12:12 h dark–light cycle with free access to standard rodent chow. Pregnant females were euthanized at GD18 (2.5% isoflurane). Plasma and tissues were collected via snap freezing. Pups and placental weights were collected ex-utero.

### Osmotic Minipump Chronic Infusion

On GD11, mice were anesthetized with isoflurane (2%) in an oxygen flow (1 L/min) and implanted with subcutaneous mini osmotic pump (1007D, Alzet, USA, 0.5ul/hour) via an interscapular transverse incision. Pumps were filled with either sFlt-1 (1.2 µg/day/7days; recombinant mouse VEGFR1/Flt-1 Fc chimera protein, R&D systems #7756-FL-050) or Allo-aca leptin receptor antagonist (1.2 µg/day/7days) or combination. Sham mice were subjected to the same surgery with subcutaneous saline control.

### Vascular Reactivity

Second-order mesenteric arteries were collected from pregnant dams at GD18 after euthanasia. Arteries were cleaned from perivascular adipose tissue and cut into 2 mm length to be mounted on 40 µm wire of a DMT® wire myograph (Ann Arbor, MI) as described (baseline tension 13.1 kPa)^30^. Vessels were challenged with acetylcholine in presence and absence of nitric oxide synthase inhibitor Nω-Nitro-L-arginine methyl ester hydrochloride (L-NAME,100μM, 20min preincubation), sodium nitroprusside and phenylephrine (1nM -30μM concentrations). Maximum response to KCl was performed (80mM). Data were recorded and analyzed using LabChart analysis software (AD Instruments®, Colorado Springs, CO) as previously described.^31–34^

### Uterine Artery Resistance Index (UARI)

At GD17, UARI was measured in pregnant mice using Doppler ultrasound (Vevo 3100, Visual Sonics, Canada).^35^ Briefly, mice were anesthetized (2% isoflurane, Oxygen, 1L/min) and placed on a heated ultrasound stage. A Doppler ultrasound recording of the uterine artery was performed and Doppler velocimetry taken at 3 separate waveforms. Peak systolic velocity (PSV), and end-diastolic velocity (EDV) was captured to measure the UARI using the following equation: UARI= (PSV-EDV)/ PSV. An average of 3 UARI measurements were calculated for each mouse.

### Euthanasia and Fetal Assessment

On GD18, mice were weighed before being euthanized. Midline incision was made to expose the uterus. Litter size and viable pup counts were counted and recorded from each uterine horn. Viable fetuses and their corresponding placentas were extracted and weighed, and placental efficiency was calculated. Tissues were quickly snap frozen in liquid nitrogen and stored at −80°C, and plasma was isolated by centrifugation.

### Statistics

Data are presented as mean ± SEM. Data were analyzed by Student’s *t*-test, one-way analysis of co-variance (ANOVA), or two-way ANOVA followed by post hoc comparison test, as appropriate, using Prism 10 or later versions (GraphPad Software, San Diego, CA). Vascular reactivity curves were analyzed by two-way ANOVA with repeated measures for dose response. Differences were considered statistically significant at P<0.05.

## Results

### VEGFR1 and VEGFR3 are the predominant VEGF receptors in human trophoblasts

We compared trophoblast cell VEGFR mRNA expressions to human umbilical venous endothelial cells (HUVECs). In Figure 1A we demonstrate that human trophoblast BeWo cells express lower VEGFR1 (*Flt-1*) and R2 (*FLK-1*) expressions than HUVEC cells, however, a significantly increased expression level of VEGFR3 *(Flt-4)*. In Figure 1B we compare all 3 receptors in BeWo cells and show that indeed, VEGFR3 levels far surpass those levels of VEGFR1 and 2, and further, that the lowest expressed VEGFR in trophoblasts is VEGFR2. In fact, Ct values of VEGFR2 expression in BeWo cells approached undetectable range (average Ct=36.5±0.34). We show in Figure 1C that this expression pattern of trophoblasts of elevated VEGFR1 and VEGFR3 and low VEGFR2 is differential to endothelial cells in which the expected pattern of VEGFR2>VEGFR1>VEGFR3 mRNA expressions presented. sFlt-1 treatment did not alter endogenous levels of VEGF and PLGF in trophoblast cells, as levels in the media were unchanged (Figure S1).

**Figure 1.**
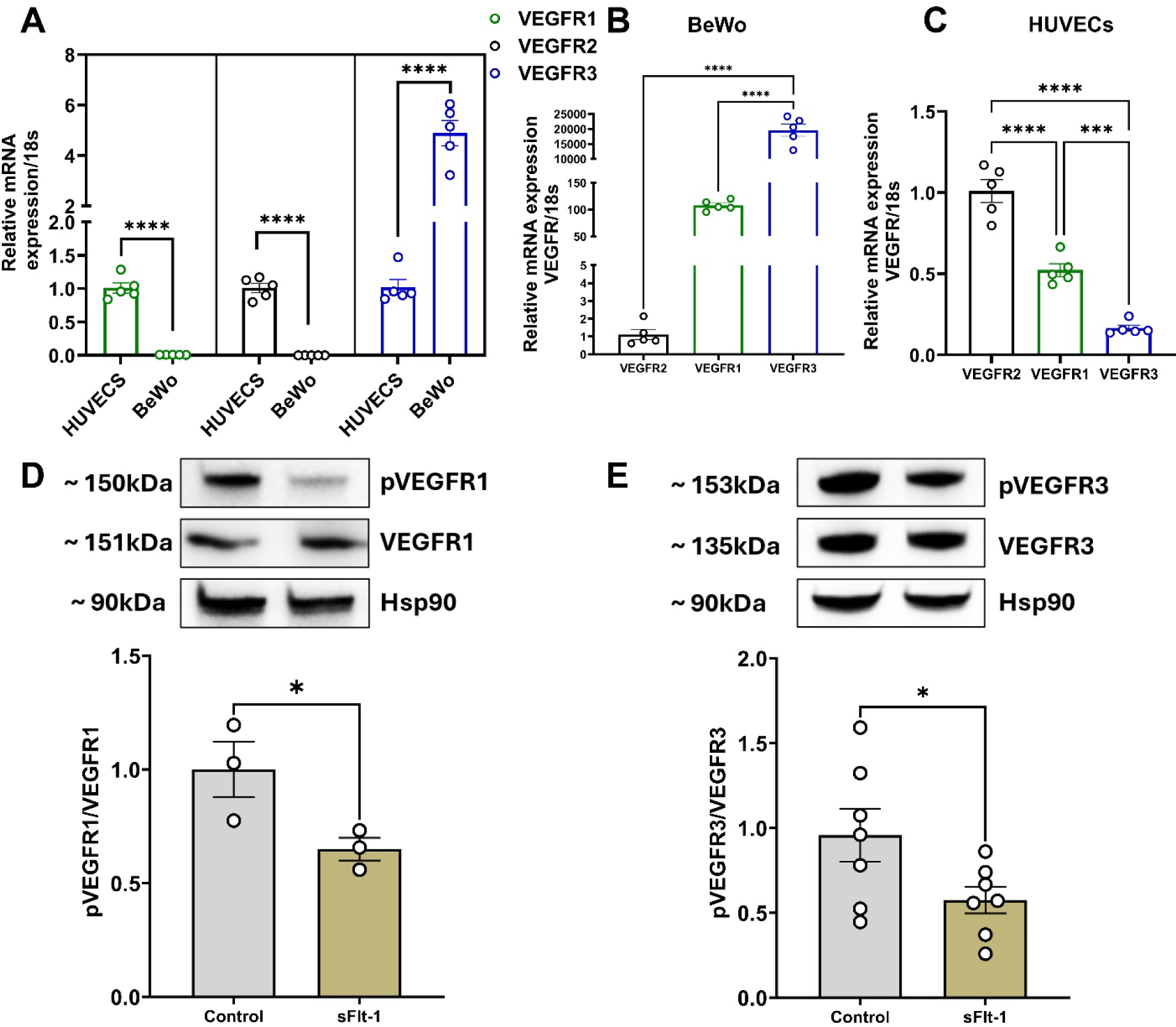
VEGFR1 and VEGFR3 are Predominant in Trophoblasts. Relative mRNA expression of VEGFR1, VEGFR2, VEGFR3 in both HUVECs and BeWo cells **(A)**; VEGFR gene expression in BeWo cells **(B)**; and HUVECs **(C)**. BeWo cells treated with either a vehicle or sFlt-1 showing the protein levels of pVEGFR1 **(D)**; and pVEGFR3 **(E)**; One Way ANOVA, Tukey’s post hoc test for A-C. Unpaired t test in D,E. *p<0.05,***p<0.001, ****p<0.0001.

### sFlt-1 decreases phosphorylation of VEGFR1 and 3 in human trophoblasts

We measured VEGFR1, 2 and 3 phosphorylation in BeWo trophoblasts and HUVECs via western blotting. We found VEGFR2 phosphorylation and total expression to be undetectable in BeWo cells (data not shown). In Figure 1D and E, VEGFR1 and 3 phosphorylation decrease with sFlt-1 incubation in BeWo cells. To confirm that our treatment of sFlt-1 is sufficient to induce classical inhibition of VEGFR signaling, we show that phosphorylation of extracellular signal-regulated kinases (ERK) increases with VEGF, and decreases with sFlt-1+VEGF in endothelial, but not in trophoblast cells (Figure S2 A and B).

### sFlt-1 treatment does not alter genes related to VEGF signaling in trophoblast cells

We performed RNA sequencing for BeWo trophoblast cells treated with sFlt-1 and vehicle. Data showed that there are 600 differentially expressed genes (DEG) between control and sFlt-1 treated cells. Figure 2A shows a heatmap featuring the top differentially expressed genes between control and sFlt-1 groups. We found 315 upregulated genes and 285 downregulated genes between control and sFlt-1 groups (Figure 2B). We performed pathway analysis to determine genes altered in sFlt-1-treated cells that were associated with VEGF signaling (Figure 2C). No genes were downregulated more than 2-fold belonging to the VEGF signaling pathway, all genes that were associated with VEGF signaling were modestly upregulated. Therefore, the VEGF signaling-related genes in trophoblasts likely differ from those that are known in other cell types and featured in pathway analysis.

**Figure 2.**
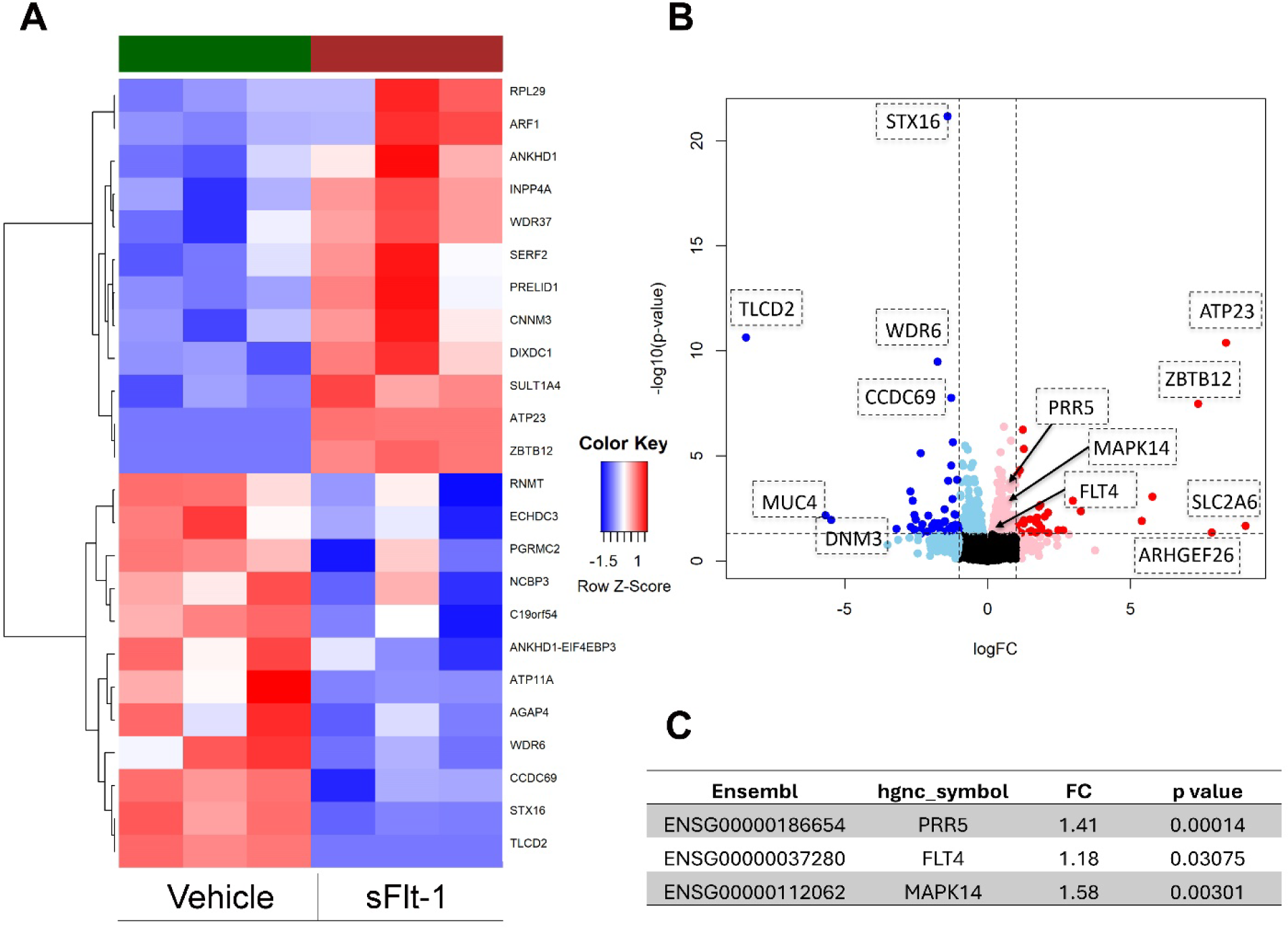
RNA sequencing of BeWo trophoblasts with and without sFlt-1 treatment. Heatmap of top DEG in response to sFlt-1 in BeWo cells **(A)**; Volcano plot showing the upregulated (red) and downregulated (blue) genes **(B)**; DEG related to VEGF signaling pathways **(C)**.

### sFlt-1 induces leptin secretion in human placental explants and human trophoblasts

We performed immunostaining for BeWo human trophoblast cells against leptin after incubation with either sFlt-1 or vehicle and imaged via confocal microscopy (Figure 3A,B). Trophoblast cells elevate leptin staining both intracellularly and in the extracellular space when treated with sFlt-1. In Figures 3C-E we measured leptin production in media of trophoblast cells and placental explants from preeclampsia and healthy patients. We show that in trophoblast cells, as well as preeclampsia and healthy pregnant placenta, sFlt-1 significantly increases leptin concentration in culture media. Neither VEGF nor PlGF alone significantly altered trophoblast leptin secretion in cells or explants, however, addition of VEGF, PLGF or VEGF+PLGF to culture media ablated the ability of sFlt-1 to increase leptin concentration in media. In preeclampsia explant cohorts, combination treatment of VEGF+PlGF had the most robust effect to suppress sFlt-1-mediated increase in leptin production. Therefore, sFlt-1 is a novel driver of increases in placental leptin production in placental trophoblasts, which is ablated by additional treatment of cells with VEGF or PlGF.

**Figure 3.**
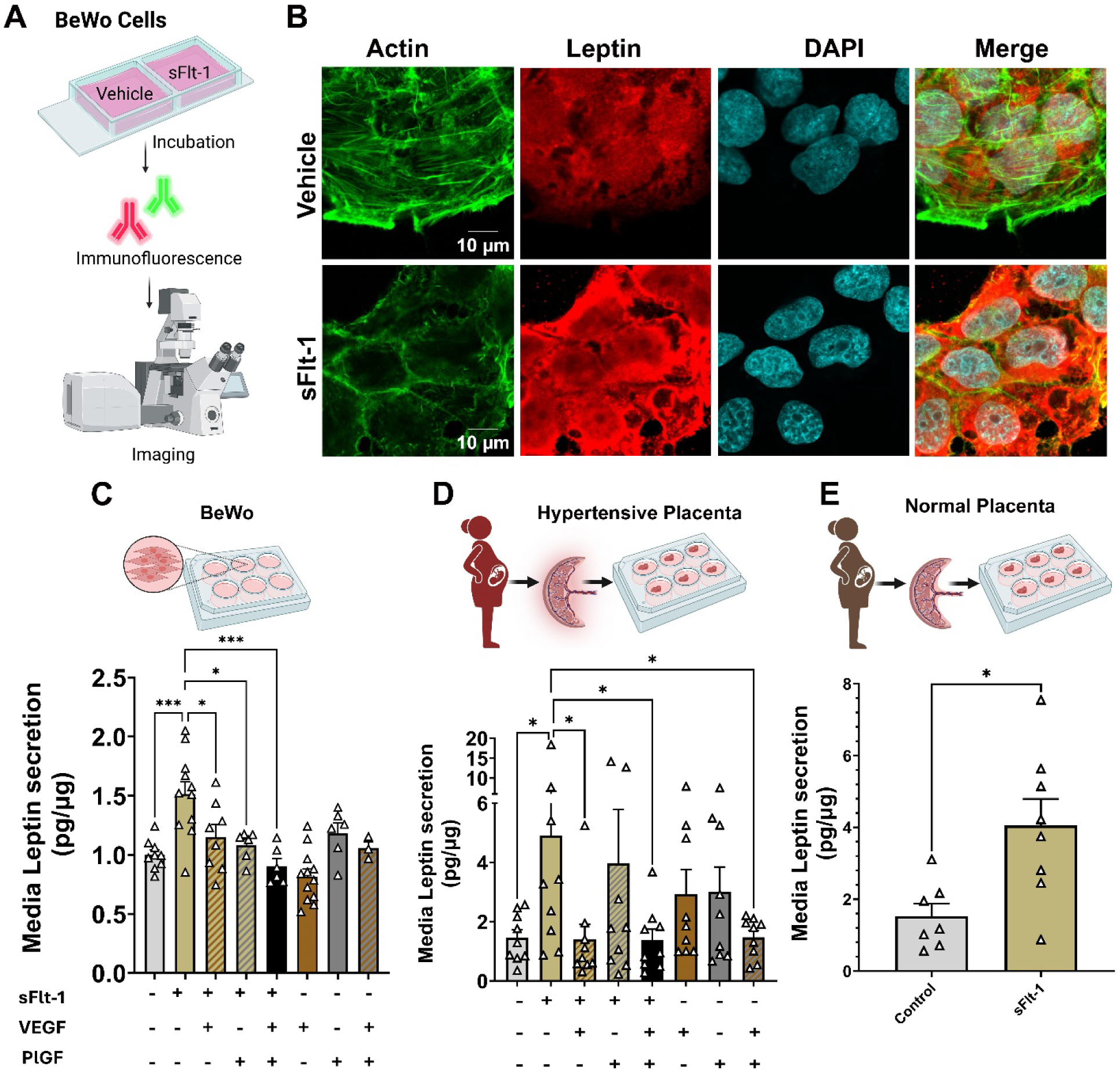
sFlt-1 induces leptin production in human placenta, which is ablated by VEGF and PlGF. Schematic showing immunostaining experiment design (**A)**; immunohistochemistry for BeWo cells treated with a vehicle (top row) or sFlt-1 (bottom row) for 24 hours (**B**); Leptin secretion in media after treatment with vehicle, sFlt-1 (72 hours), VEGF (24 hours), PlGF (24 hours) or combination in Bewo cells **(C)**; placental explants from preeclampsia pregnancies **(D)** or placental explants from normal pregnancies **(E);** One Way ANOVA, Fisher’s LSD post hoc test, Student’s t-test. *p<0.05, ***p<0.001.

### Silencing of either VEGFR1 or 3 upregulates trophoblast leptin production

We performed VEGFR1, VEGFR2 and VEGFR3 gene silencing in BeWo cells using siRNA. We show that silencing effectively reduced expression of respective VEGFR expressions, except for VEGFR2 in which we were unable to detect expression both with and without silencing (data not shown) (Figure 4A,B). In Figure 4C, we show that VEGFR1 siRNA, VEGFR3 siRNA and VEGFR1+3 siRNA, all individually had a significant effect to increase leptin concentration in culture media, when compared to both control and sFlt-1 treatment. As expected with the lack of expression observed we saw that VEGFR2 silencing did not alter leptin secretion from trophoblast cells (Figure S3). Trophoblast cells were treated with VEGF_165_b, a VEGFR1/2 agonist, and VEGFC, a VEGFR2/3 agonist. Treatment with either VEGF_165_b or VEGFC resulted in a significant blunting of the ability of sFlt-1 to increase media leptin production (Figure 3F). Therefore, inhibition of VEGFR1 and 3 signaling downregulates leptin production in trophoblasts.

**Figure 4.**
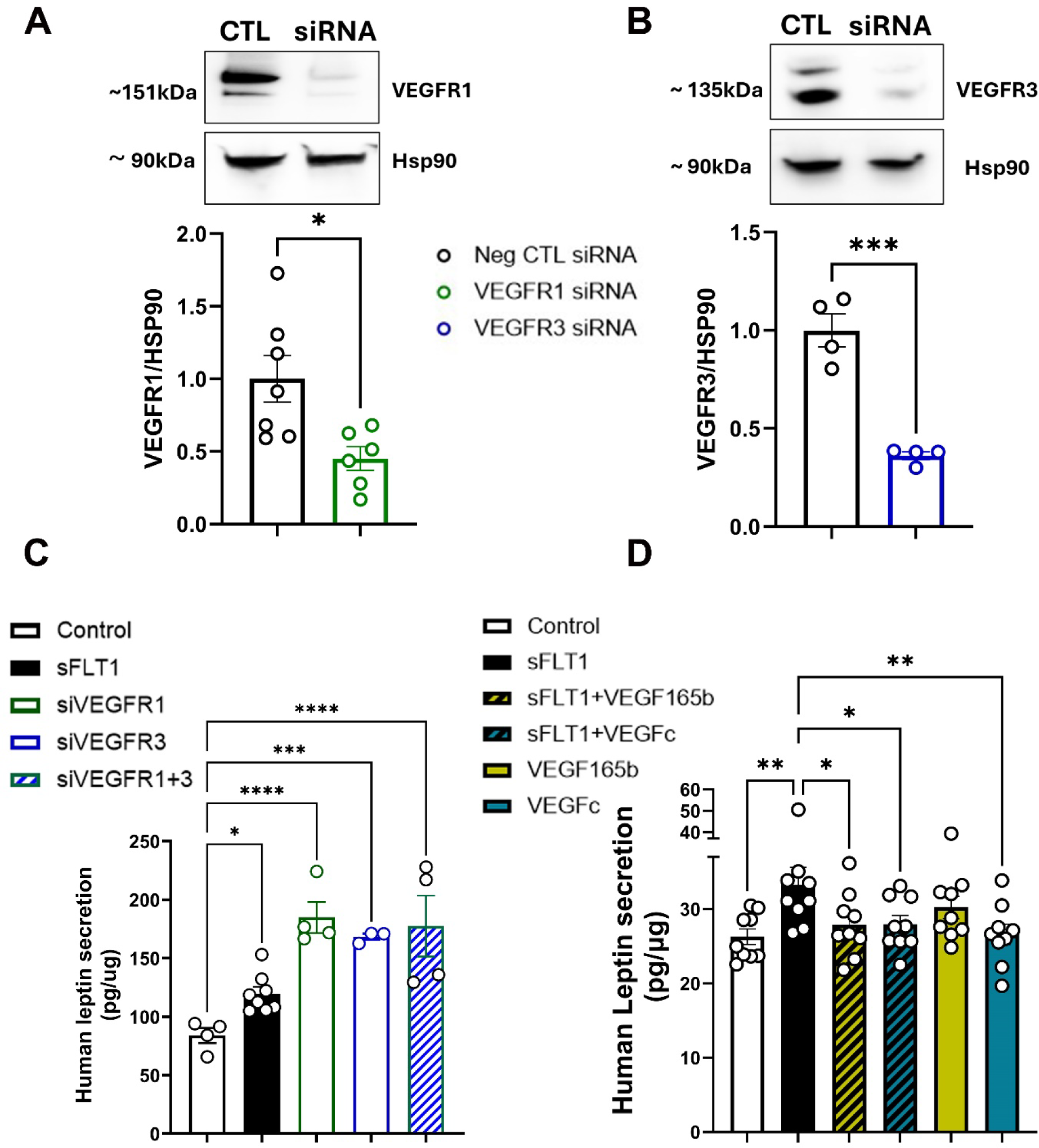
VEGFR1 and VEGFR3 induce leptin production in trophoblasts. Western blot showing the effect of VEGFR1 siRNA **(A)** and VEGFR3 siRNA **(B)** on their respective receptor protein expression. Leptin secretion level measured in the media of BeWo cells treated with sFlt-1, siVEGFR1, siVEGFR3, siVEGFR1+ siVEGFR3, or a vehicle **(C)**; Leptin secretion level measured in the media of BeWo cells treated with sFlt-1, VEGF165b, VEGFc, vehicle or a combination **(D)**. Unpaired t-test in figure 3A-D. One Way ANOVA, Fisher’s LSD post hoc test for 3E-F. *p<0.05, **p<0.01, ***p<0.001, ****p<0.0001.

### sFlt-1 increases plasma leptin levels in pregnant mice

We infused mice with sFlt-1 from GD11-18. In Figure 5A we show that sFlt-1 significantly increases plasma leptin levels in pregnant mice. In contrast to higher order mammals such as humans and primates, mice and rats produce nondetectable or minimal leptin in placental trophoblasts.^36,37^ To assess whether sFlt-1 induced an upregulation of leptin, we measured mRNA transcript in subcutaneous adipose tissue biopsies from Balb/C pregnant mice at GD11, 15 and 18. At GD11 prior to sham/sFlt-1 infusion leptin mRNA expression did not differ, however, sFlt-1 infusion induced a significant upregulation of leptin mRNA at GD18 in pregnant mice (Figure 5B). Therefore, similar to human placental trophoblasts, sFlt-1 induces upregulation of adipose leptin production and leptin levels in intact *in vivo* rodent pregnancy.

**Figure 5.**
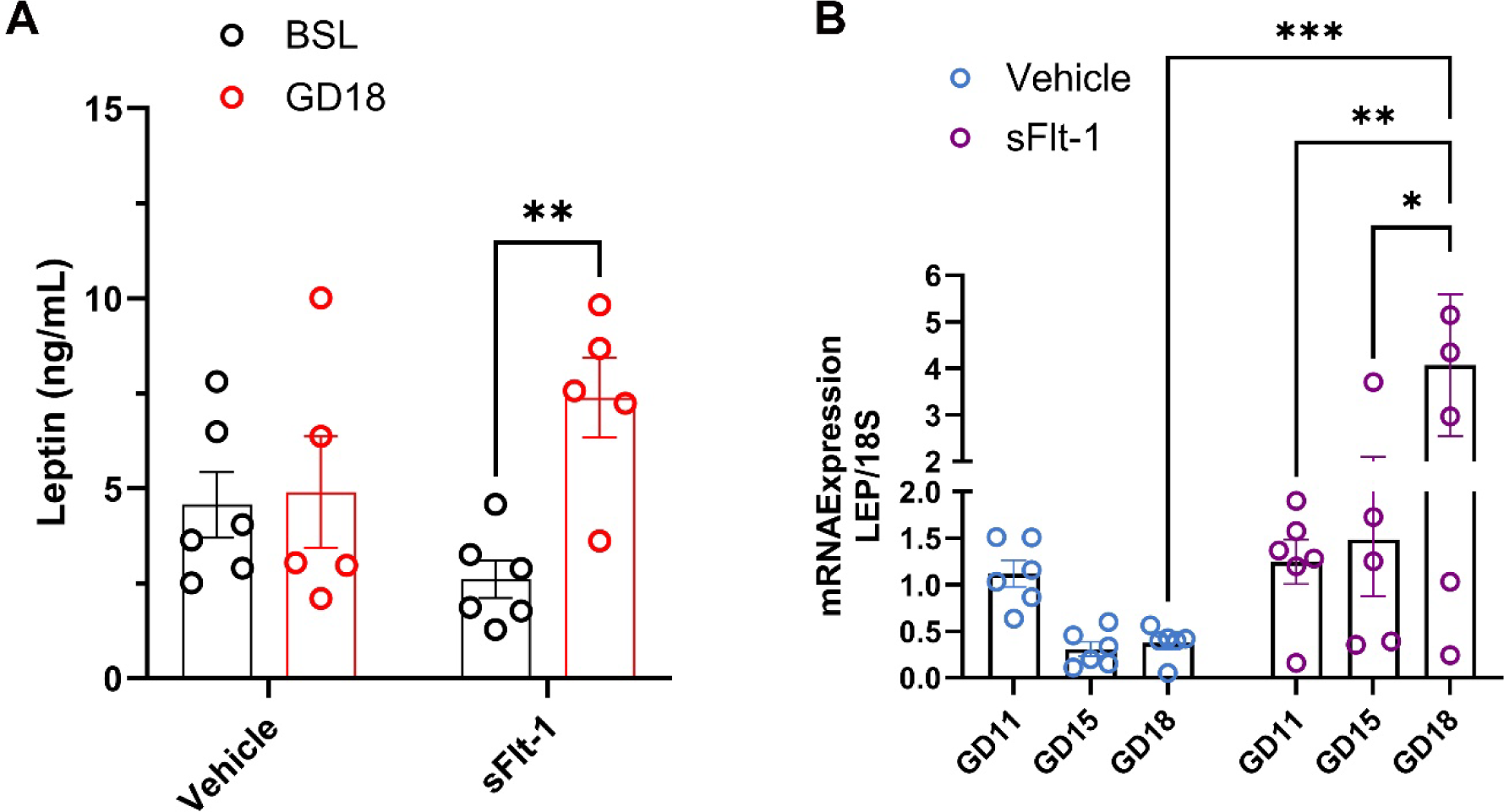
Leptin plasma levels and gene expression in sFlt-1-treated pregnant mice. Plasma leptin levels collected at baseline and GD18 for mice treated with sFlt-1 or a vehicle **(A);** Leptin gene expression in adipose tissue samples collected at GD11, GD15 and GD18 in mice treated with sFlt-1 or a vehicle **(B).** Two-way ANOVA, Fisher’s LSD post hoc test. *p<0.05, **p<0.01, ***p<0.001.

### sFlt-1 induces vascular dysfunction in pregnant mice, which is ablated by leptin receptor antagonism

Clinical reports demonstrate that high sFlt-1 is a predictor of adverse outcomes in preeclampsia patients^38^ and several studies indicate that sFlt-1 induces characteristics of preeclampsia in pregnant rodents.^39,40^ However, whether sFlt-1 infusion alone induces vascular endothelial dysfunction in experimental models has yet to be demonstrated. Furthermore, we recently demonstrated that elevated leptin levels is a crucial mediator of endothelial dysfunction in mouse pregnancy.^26,41^ We infused mice with sFlt-1 alone or in combination with leptin receptor antagonist (allo-aca) from GD11-18 via subcutaneous osmotic minipump (Figure 6A). At GD17 sFlt-1 increased UARI in pregnant mice compared to sham females (Figure 6B). Allo-aca alone did not affect UARI in pregnant mice and further had no effect to reduce UARI in sFlt-1-infused pregnant mice. In isolated mesenteric arteries sFlt-1 reduced endothelial-dependent relaxation to acetylcholine (ACh) assessed as the dose-response (Figure 6C), the maximal response and the EC_50_ (Table 1). The effect of sFlt-1 to reduce ACh-mediated relaxation was blunted in mice treated with allo-aca in all metrics assessed (Figure 6C and Table 1). Incubation with nitric oxide synthase (NOS) inhibitor N(G)-Nitro-L-arginine methyl ester (L-NAME) abolished any differences between groups in ACh-mediated relaxation, suggesting that decreases in NO synthesis drives sFlt-1-mediated endothelial dysfunction (Figure 6D). Neither sFlt-1 or allo-aca altered vascular contractility to α-1 agonist phenylephrine (Phe, Figure 6E, Table 1). sFlt-1 significantly reduced endothelial-independent relaxation in response to sodium nitroprusside (SNP, Figure 6F, Table 1). Allo-aca ablated the ability of sFlt-1 to reduce SNP-mediated relaxation in mesenteric vessels. Therefore, leptin receptor activation is required for sFlt-1-mediated vascular dysfunction, but not necessarily adaptations to uterine artery flow.

**Figure 6.**
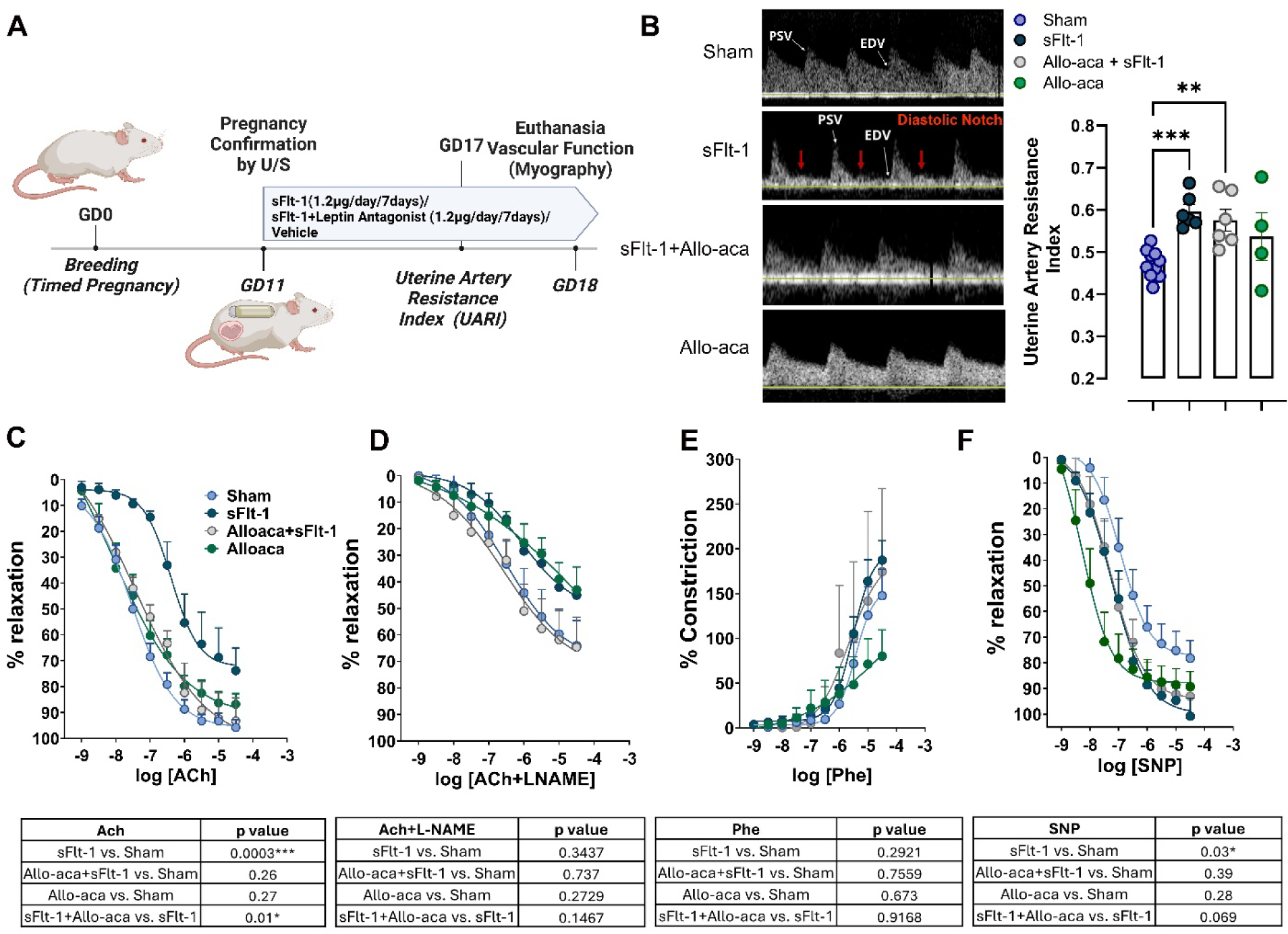
sFlt-1 induces vascular dysfunction dysfunction, which is reversed by leptin antagonist. A schematic for mouse experimental timeline **(A)**; Uterine artery resistance index measured using Doppler ultrasound **(B)**; Vascular reactivity measured in second-order mesenteric arteries of different experimental groups in response to Ach **(C)**, Ach+L-NAME **(D)**, Phe **(E)**, SNP **(F)**. Two-way ANOVA with and without repeated measures. *p<0.05, **p<0.01, ***p<0.001.

**Table 1.** Relaxation and contractile responses to ACh, Phe and SNP expressed as the maximal response at the highest dose (“Max”) and the half maximal effective concentration (EC50).

|  | **Sham** | **sFlt-1** | **Alloaca+sFlt-1** | **Alloaca** |
| --- | --- | --- | --- | --- |
| Ach Max | 95.75± 2.883 | 73.83±8.731* | 93.2±8.885 | 86.75±4.029 |
| Phe Max | 147.7±29.55 | 187.3±21.78 | 174.5±92.75 | 80.08±29.37 |
| SNP Max | 89.25±5.808 | 78.17±6.863 | 98±6.595<sup>#</sup> | 93.6±2.315 |
| Ach EC<sub>50</sub> | -7.567±0.1601* | -5.660±0.6006<sup>††</sup> | -7.368±0.3466 | -9.072±1.731 |
| Phe EC<sub>50</sub> | -5.380±0.1057 | -5.471±0.1108 | -2.431±2.370 | -5.687±0.4419 |
| SNP EC<sub>50</sub> | -8.250±0.2524* | -6.892±0.1317 | -6.986±0.2321 | -7.298±0.5108 |

Maximal relaxation is percentage of precontraction
\*p<0.05 vs sFlt-1
Maximal constriction is percentage of KCl constriction
<sup>#</sup>p<0.05 vs Alloaca+sFlt-1
EC<sub>50</sub> is the log transformed EC<sub>50</sub>
<sup>†</sup>p<0.05 vs Alloaca
All data expressed as mean ± standard error

In Figure S4 we show that, similar to other studies of sFlt-1 infusion into pregnant mice, sFlt-1 infusion alone from GD11-18 does not significantly reduce pup weight or increase fetal demise in pregnant mice. Importantly, allo-aca does not decrease fetal growth nor fetal death rate in pregnant mice in either group administered, and rather increases fetal growth. Therefore, leptin receptor antagonism does not appear to have detrimental effects on fetal programming or growth in pregnancy or in the sFlt-1-induced preeclampsia rodent model.

## Discussion

Numerous clinical cohorts across over two decades show that preeclampsia patients present with elevations in plasma sFlt-1 levels^42–45^ and this soluble form of the VEGFR1 has become a hallmark of the disease. Lesser known is that decades of patient studies also demonstrate elevations in plasma leptin levels in preeclampsia, independent of their body mass index.^14,15,46–53^ This study provides key mechanistic insight into how trophoblast-specific VEGF receptor signaling is altered by sFlt-1 to trigger increases in leptin production.

The placenta is a fleeting physiological organ, developing alongside the growing fetus, regulating all aspects of its growth and nutrition while secreting hormones that trigger adaptations in maternal organ systems. The placental trophoblasts are the predominant cell type in the placenta and perform numerous processes throughout gestation, however, the regulation of their various functions remains underdeveloped, particularly in disease states such as preeclampsia. Trophoblast invasion of the uterine muscle tissue is a key process that initiates spiral artery remodeling in pregnant women, facilitating the high vascular capacity needed for a growing fetus.^54^ In accordance, heightened angiogenesis is needed to elongate and expand these arteries and increase endothelial proliferation, which is provided by systemic elevations in VEGF and PlGF production and VEGFR activation.^55,56^ It has long been known that the trophoblasts themselves express both the soluble (sFlt-1) and membrane-bound forms of VEGFR1 (Flt-1)^57^ and VEGFR3 (Flt-4),^58^ but the function of these receptors in this cell type is largely unexplored.

In endothelial cells, VEGFR2 is the predominant receptor primarily mediating angiogenic activity in early pregnancy. VEGFR1 is largely thought to act as a “decoy” receptor in endothelial cells though recent papers suggest VEGFR1 can signal through STAT3 phosphorylation and other down-stream candidates have been proposed.^10,59–61^ VEGFR3 on the other hand is predominantly expressed in lymphatic epithelial cells, with little known function in other cell types.^62^ We sought to determine which VEGF receptor subtypes are predominant in trophoblastic cells and whose activity are impacted by elevated sFlt-1. Unlike endothelial cells, our data show that VEGFR2 is minimally expressed, undetectable by western blot, whereas both VEGFR1 and VEGFR3 are highly expressed in human trophoblasts. These findings indicate that VEGFR1 and 3 are not likely decoy receptors in trophoblast cells, but rather induce major signaling cascades that result in protein synthesis.^63^ Importantly VEGFR1 and R3 have unique ligands, R1 binding both VEGF and PlGF, while only VEGF will activate R3. In our study we show that sFlt-1 in our *in vitro* trophoblast environment reduces phosphorylation of both R1 and R3, while not affecting the levels of VEGF/PlGF in the media. Therefore, sFlt-1 is reducing activity of R1 and R3, but not likely influencing production of ligand.

Angiogenic imbalance in preeclampsia is a well-established notion, and the FDA recently approved an sFlt-1/PLGF ratio as a prognostic test for onset of preeclampsia severe features to guide care strategies.^64,65^ sFlt-1 is a truncated form of VEGFR1 that sequesters VEGF and PlGF in the extracellular space.^10^ In this manner, sFlt-1 acts as an anti-angiogenic factor, given that VEGF and PlGF are the primary ligands of the membrane-bound VEGFRs. Many studies show impaired development of placental spiral artery vasculature in patients who will go on to develop preeclampsia, and it may fall to reason that sFlt-1 promotes an anti-angiogenic state that restricts the spiral arteries, ultimately leading to preeclampsia. However, cohorts show that elevations in sFlt-1 are not consistently early enough in gestation to affect spiral artery remodeling ^66,67^, and also, that spiral artery remodeling deficits are variably found in preeclampsia cohorts.^68,69^ Therefore, sFlt-1 likely promotes the development of preeclampsia via mechanisms beyond simply inhibiting angiogenesis of placental vasculature. In this study, we propose a novel mechanism for sFlt-1-mediated vascular damage in its ability to significantly upregulate leptin production.

Leptin is peptide hormone most commonly associated with obesity, and in nonpregnant individuals adipose tissue is overwhelmingly the main source of leptin production. However, in pregnancy, placental trophoblasts become a highly potent source of leptin production. Although a rise in plasma leptin levels across pregnancy is both normal and required for placentation^70^, preeclamptic patients present with distinctly elevated plasma leptin levels and trophoblast leptin production, independent of body mass index.^14,15,46–53^ Our study shows for the first time a potential mechanism whereby sFlt-1 may lead to this increase in leptin production in preeclampsia patients. In both freshly collected human placental explants from preeclampsia patients and human BeWo trophoblast cells we show that sFlt-1 elevates leptin secretion into culture media, and that either VEGF or PlGF blocks this effect. As the levels of VEGF/PlGF in media do not change with sFlt-1 treatment, it is likely that sFlt-1 sequestration of these two peptides leads to upregulation of leptin production in trophoblasts. We show that VEGFR1 or VEGFR3 silencing, without addition of sFlt-1, potently upregulates leptin production in trophoblast cells, beyond that of sFlt-1 alone. And further, we show that treatment with the VEGFR1-specific ligand VEGF_165b_ and the VEGFR3-specific ligand VEGFC are able to ablate sFlt-1-mediated increases in leptin production in trophoblasts. We, therefore, propose that placental trophoblasts produce leptin in a steady state which is regulated by VEGFR1 and R3 signaling, and the downregulation of this signaling triggered by sFlt-1 removes a molecular “brake” on leptin production.

Our data suggest that VEGFR1 and R3 may trigger a similar intracellular signaling cascade that leads to suppression of leptin production, warranting further study into the signaling of these receptors in trophoblasts. We show evidence that intracellular signaling of VEGFR1 and R3 in trophoblasts is not similar to that of VEGFR2 in endothelial cells. We demonstrate that MAP kinase pathway activation, via phosphorylation of ERK1/2, is not altered by sFlt-1 in trophoblasts as it is in HUVECs. We additionally performed an RNA sequencing of BeWo cells treated with sFlt-1 and found that although a small number of genes were up- and down-regulated, no genes associated with angiogenesis were found to be significantly downregulated >2-fold. In fact, the 3 genes in this pathway identified with any change (*PRR5, Flt-4* and *MAPK14*) were modestly upregulated at very low fold changes. Therefore, trophoblast VEGFR signaling is likely unique to what is currently in the literature for other cell types and other as-yet undiscovered intracellular signaling cascades are likely at play that decrease leptin production upon activation of VEGFR1 and R3, which warrant future study.

Preeclampsia is a disease of placenta origin, however, growing evidence shows that vascular endothelial dysfunction is a hallmark of the disorder and precedes clinical features.^46,65,71^ We recently showed that high leptin levels in murine pregnancy are essential for vascular endothelial dysfunction in both normal pregnant mice and mice with placental ischemia.^26,41^ Leptin is a cardiovascular regulator that induces vascular endothelial dysfunction and mediates adverse cardiovascular outcomes pronouncedly in females.^31–33^ We show novel data that sFlt-1 infusion in mid-late gestation of pregnant mice induces vascular endothelial and smooth muscle dysfunction in the resistance arteries in late pregnancy. We additionally show that vascular resistance at the uterine artery is increased by sFlt-1. Although we have shown that leptin administration to pregnant rodents induces increases in blood pressure^72^, administration of a leptin receptor antagonist *in vivo* in pregnancy has not been reported to-date. Our data suggest that allo-aca leptin receptor antagonist is able to ablate the effect of sFlt-1 when administered mid-late gestation on vascular function in both the endothelial cells and vascular smooth muscle cells. We did observe that allo-aca did not improve uterine artery resistance index, indicating that sFlt-1 may have direct non-leptin mediated placental outcomes that prevented improvement of flow velocity across the feto-placental interface. Importantly, however, allo-aca did not reduce fetal growth, and increased growth compared to sham. Recent studies from our laboratory suggest that leptin is a key mediator of placental mitochondrial protein unfolding responses and lipid accumulation.^72^ Therefore, by inhibiting leptin receptors in placenta mitochondrial function may have increased to promote greater nutrient exchange.

There are a few limitations to the current study. The placental explant studies were all done in term placentas, therefore, earlier gestation events, such as those that occur in early placentation, were not developing at the time of these experiments to test trophoblast endocrine function. Further, the sFlt-1 model of preeclampsia utilized (mid-gestation infusion), like all models of pregnancy disorders in rodents, is limited by the vast differences in murine pregnancy compared to humans. Rodent placenta do not produce leptin in levels comparable to humans and large-order mammals with primarily singleton pregnancies.^36,37,73^ Therefore, the effects on maternal function in large order mammals with intact trophoblast leptin production can only be speculated from this study. However, the endothelial dysfunction phenotype observed in our mouse model is comparable to the presentation of systemic endothelial dysfunction in preeclampsia patients, therefore, we believe that this report significantly moves the field forward with identifying key mechanisms whereby the endocrine disruption of the preeclamptic placenta results in systemic cardiovascular consequences.

## Conclusions and Perspectives

The present study presents key findings that uncover a previously unrecognized VEGF-mediated endocrine disruption in a non-endothelial cell type, the trophoblast cell. We identify for the first time the expression levels of the VEGF membrane receptors in human trophoblasts compared to endothelial cells and identify a novel role for VEGFR1 and 3 as molecular suppressors of trophoblast leptin production. We additionally provide the first direct evidence that sFlt-1, and subsequent disruption of trophoblast VEGFR1 and 3 phosphorylation, is the likely mediator of increases in plasma leptin levels that have been long observed in preeclampsia patients. These findings lay important groundwork for future investigations into targeted inhibition of this pathway, with the potential to inform novel treatment strategies for the severe cardiovascular and fetal complications associated with preeclampsia.

## Acknowledgments

None

